# Chronic clinical signs of upper respiratory tract disease shape gut and respiratory microbiomes in cohabitating domestic felines

**DOI:** 10.1101/2022.05.09.491187

**Authors:** Holly K. Arnold, Rhea Hanselmann, Sarah M. Duke, Thomas J. Sharpton, Brianna R. Beechler

**Author notes:** Indicates these authors contributed equally to this work and are listed alphabetically (Thomas J. Sharpton, Brianna R. Beechler).

## Abstract

<u>F</u>eline <u>u</u>pper respiratory tract <u>d</u>isease (FURTD), often caused by infections etiologies, is a multifactorial syndrome affecting feline populations worldwide. Because of its highly transmissible nature, infectious FURTD is most prevalent anywhere cats are housed in groups such as animal shelters, and is associated with negative consequences such as decreasing adoption rates, intensifying care costs, and increasing euthanasia rates. Understanding the etiology and pathophysiology of FURTD is thus essential to best mitigate the negative consequences of this disease. Clinical signs of FURTD include acute respiratory disease, with a small fraction of cats developing chronic sequelae. It is thought that nasal mucosal microbiome changes play an active role in the development of acute clinical signs, but it remains unknown if the microbiome may play a role in the development and progression of chronic clinical disease. To address the knowledge gap surrounding how microbiomes link to chronic FURTD, we asked if microbial community structure of upper respiratory and gut microbiomes differed between cats with chronic FURTD signs and clinically normal cats. We selected 8 households with at least one cat exhibiting chronic clinical FURTD, and simultaneously collected samples from cohabitating clinically normal cats. Microbial community structure was assessed via 16S rDNA sequencing of both gut and nasal microbiome communities. Using a previously described ecophylogenetic method, we identified 37 and 27 microbial lineages within gut and nasal microbiomes respectively that significantly associated with presence of active FURTD clinical signs in cats with a history of chronic signs. Overall, we find that nasal and gut microbial communities may contribute to the development of chronic clinical course, but more research is needed to confirm our observations.

## Introduction

<u>F</u>eline <u>u</u>pper respiratory tract <u>d</u>isease (FURTD), most often caused by infectious etiologies, is a highly prevalent multifactorial syndrome that affects felid populations worldwide. Due to the highly contagious nature of infectious FURTD pathogens, infectious FURTD is most prevalent anywhere cats live in groups (1), and thus FURTD is a major burden on animal shelters (2). Clinical signs of infectious FURTD include ocular or nasal discharge, sneezing, epistaxis, and severe ocular, nasal, or oral pathology including inflammation and ulceration of mucous membranes and corneas (3, 4). Even though morbidity greatly surpasses mortality in natural infection course (5), infectious FURTD can increase euthanasia rates in shelters (6) by increasing time to adoption, contributing to overcrowding, and shunting resources into costly clinical treatments, and isolation/sanitization protocols (5). Understanding the etiology of infectious FURTD is a thus priority in feline medicine to better mitigate the negative consequences of the disease.

Unfortunately, both FURTD etiology and pathophysiology remain difficult to unravel in part because of the complexity of the disease. At least six primary pathogens have been identified as causing signs of disease, including viral (Feline Herpes Virus 1, Feline Calicivirus), bacterial (*Bordetella bronchiseptica*, *Chlamydophila felis*, *Mycoplasma species*), and less commonly fungal (*Cryptococcus neoformans*) organisms. Infection with a primary agent is thought to proceed a secondary change in underlying microbial community composition, both of which likely contribute to clinical signs. Using culture-independent 16S sequencing, prior work has shown that nasal microbial communities are indeed shaped by the presence of primary FURTD pathogens during acute infection (7). Some, but not all, cats will go on to develop a chronic disease course from presence of latent infection (i.e. Feline Herpes Virus 1), or an immune mediated response to infection (5). It remains unknown, however, if secondary microbial changes persist in cats with chronic clinical disease and if such changes contribute to persistence of clinical signs.

We reasoned that cats with long term FURTD signs may develop a chronicity in clinical course due to unique interactions between primary FURTD pathogen and persistent changes within the respiratory microbiome. For example, widely reported detection ranges (9 - 89%) for Feline Herpes Virus 1 (FeHV-1) in cats with clinical signs consistent with FURTD (8–17), as well as identification of FeHV-1 in cats suspected to be clinically normal (3 - 49%) (8,12,14,15,17,18), begs the question on why some cats with primary etiological agents do not chronically present with clinical signs while others do. We hypothesized that cats identified as having chronic clinical manifestations consistent with infectious FURTD signs may have upper respiratory microbiome changes compared to cats without chronic clinical signs.

Recent studies increasingly note the presence of a gut-respiratory axis, resulting in a dynamic interplay between gut and airway microbial communities modulated by shared mucosal immunity (19–21). Changes within the gut microbiome may happen secondarily to upper respiratory tract disease as a result of systemic inflammation (22), or may be a primary cause leading to signs of upper respiratory tract disease. For example, exposure to dust microbial communities altered gut microbiome community structure in a mouse model, and consequently protected host from airway viral infection (23). Disrupted gut communities can influence immune responses at distal sites (24, 25), and microbial diversity exposure has been inversely correlated to allergy, asthma and allergic rhinitis development in humans (26–28). Therefore, we hypothesized that cats with chronic FURTD clinical signs may also have changes that are apparent within their gut microbiome.

Culture-independent 16S rDNA sequencing of the cat upper respiratory microbiome has revealed a rich and variable respiratory community, that was incompletely documented by culture techniques alone (7). Significant variability in upper respiratory microbiome composition was observed both within and among cats with and without clinical signs, as well as across age classes, sex, and breed groups (7, 29). The high variation of respiratory compositions in prior work resulted in difficulty distinguishing key microbial taxa which consistently differed between control and FURTD subjects (7), some of which may have been driven by differing environmental factors (29, 30). We reasoned that a large contributing factor to nasal and gut microbiome communities is the metacommunity to which the cat is exposed (i.e. household). Thus, we developed a study design where each household was recruited on the basis of having at least one cat with a history of chronic upper respiratory tract signs, as well as also sampling apparently healthy cats in the same household when possible.

Our objective was to further characterize the etiology of FURTD in cats with and without chronic clinical signs. We show that inflammatory markers significantly differ in cats with FURTD signs compared to those without. We then apply a novel ecophylogenetics technique (31, 32) in order to discover monophyletic microbial clades and individual microbial biomarkers that exhibit differential ecological distributions across cats based on disease status.

## Methods

### Population Recruitment & Study Design

We selected eight households where each household was recruited on the basis of having at least one cat with a history of chronic upper respiratory tract signs of at least one year duration, and also sampled apparently healthy cats in the same household if present. Clients were recruited from Oregon State University in Corvallis, Oregon, as well as the or from community members within the broader Corvallis area. All participating cats were housed primarily indoors and had proof of current rabies vaccination. Cats were required to have lived in the same household for at least one year to account for environment associated variation in microbiome communities (29). A thorough clinical history and physical examination performed by a veterinarian, and baseline diagnostics were obtained to exclude cats with concurrent disease. Cats with a history of recent (i.e. < 3 months) vaccination with modified live organisms (parenteral or intranasal), treatment with antibiotics, anti-inflammatories, or immunosuppressive drugs, or concurrent non-respiratory illness were excluded.

### Quantification of clinical signs

All cats were examined by a veterinarian to confirm the presence or absence of clinical signs. On physical exam, each clinical sign was encoded as a one if a sign of infectious FURTD and a zero otherwise for each organ system (i.e. ocular, nasal, oral, pulmonary). This resulted in matrix of systems by subject where every entry was scored as a “one” if abnormal and consistent with FURTD and a “zero” otherwise. For example, “dental tarter” was entered as a zero, because, while an abnormal physical finding, it is not a finding consistent with FURTD. Cats were classified into the “case” category if they had at least one sign consistent with chronic FURTD, and the “control” category if they did not. Clinical signs that were encoded as consistent with FURTD included ocular (bilateral serous or mucopurulent ocular discharge, conjunctivitis, chemosis, keratitis, corneal ulceration and anterior uveitis), nasal (bilateral serous to mucopurulent nasal discharge, sneezing, and rhinitis) and oropharyngeal (stomatitis, glossitis, faucitis) clinical signs.

### Classification of cases and controls

Cats were classified into cases if they had a history of chronic FURTD signs and/or the presence of at least one clinical sign consistent with FURTD was observed on veterinary physical examination (**Table 1**). Cats were classified as controls if they did not meet these criteria. PCR testing for primary FURTD pathogens were positive in four of the cases and zero of controls. Gut and nasal microbiomes were sequenced on available samples. One cat was excluded because it was unknown when the last possible antibiotic treatment occurred (Supplementary Table 1). Overall, nasal samples for nasal microbiome sequencing were available in 17 (11 cases, 6 controls). In two cats we were unable to obtain a fecal sample, and so gut microbiome sequencing was performed in only 15 individuals (10 cases, 5 controls).

**Table 1:** Study Population. Cats were recruited based on a clinical history of the presence of chronic (>1 year duration) clinical signs (n = 12). When available, cohabitating housemates defined as living in the same living situation for greater than 1 year (n = 6) were also included, for a total of 18 individuals. For each organ system (i.e. ocular, nasal, oral, pulmonary) clinical signs consistent was FURTD were encoded as a “one” if present and a “zero” if absent. Abnormal signs that were reported by a veterinarian that were included as a sign of FURTD included ocular discharge, ocular crusts, blepharospasm, nasal discharge, faucitis, and upper airway noise. A description of table columns follow. *ID:* Cat ID number. *Age:* in years*. Sex:* FS = Female Spayed; MN = Male Neutered. *Ocular/Nasal/Oral/Resp:* Signs present consistent with FURTD and confirmed by veterinary physical exam. *Status:* Case (n =12); Control (n = 6). *MB:* N = samples collected for nasal microbiome sequencing. G = samples collected for gut microbiome sequencing. * = sample excluded based on exclusion criteria. *PCR Result:* FeHV-1 = Feline Herpes Virus Type 1, FCV = Calicivirus, *Bb* = *Bordetella bronchiseptica*, *Mf* = *Mycoplasma felis*.

| ID | Household | Age | Sex | Ocular | Nasal | Oral | Resp | Status | MB | PCR Result |
| --- | --- | --- | --- | --- | --- | --- | --- | --- | --- | --- |
| 1 | A | 9 | MN | 1 | 1 | 0 | 0 | Case | N | Negative |
| 2 | A | 4 | MN | 0 | 0 | 0 | 0 | Control | G, N | Negative |
| 3 | B | 11 | FS | 0 | 1 | 0 | 0 | Case | G, N | FeHV-1 |
| 4 | B | 11 | MN | 0 | 0 | 0 | 0 | Control | G, N | Negative |
| 5 | C | 4 | MN | 0 | 0 | 0 | 0 | Control | N | Negative |
| 6 | C | 10 | FS | 1 | 0 | 0 | 0 | Case | G, N | Negative |
| 7 | D | 6 | MN | 1 | 1 | 1 | 1 | Case | N* | Negative |
| 8 | D | 9 | MN | 1 | 1 | 1 | 0 | Case | G, N | FCV, FeHV-1 |
| 9 | D | 6 | FS | 1 | 0 | 0 | 1 | Case | G, N | *Bb*, FeHV-1, *Mf* |
| 10 | D | 9 | MN | 1 | 0 | 1 | 0 | Case | G, N | FCV |
| 11 | E | 5 | FS | 1 | 1 | 0 | 1 | Case | G, N | Negative |
| 12 | E | 5 | MN | 0 | 0 | 0 | 0 | Control | G, N | Negative |
| 13 | F | 4 | FS | 1 | 0 | 0 | 0 | Case | G, N | Negative |
| 14 | F | 6 | MN | 0 | 0 | 0 | 0 | Control | G, N | Negative |
| 15 | G | 14 | MN | 1 | 0 | 0 | 0 | Case | G, N | Negative |
| 16 | G | 4 | MN | 1 | 1 | 0 | 0 | Case | G, N | Negative |
| 17 | H | 8 | FS | 1 | 0 | 0 | 0 | Case | G, N | Negative |
| 18 | H | 8 | FS | 0 | 0 | 0 | 0 | Control | G, N | Negative |

### Baseline Diagnostics

Baseline diagnostics included a chemistry panel, and complete blood count for all cats to identify inflammatory markers and screen for systemic disease. When samples were available, a urinary analysis was performed to evaluate renal function and to screen for presence of uropathology, and a fecal sample was examined for intestinal parasites. Serological evaluation for feline leukemia virus (FeLV) antigen, feline immunodeficiency virus (FIV) antibodies, and heartworm (*Dirofilaria immitis*) antigen was performed in all cats. The presence of obvious comorbidities was not found in any of the cats aside from one with lab results consistent early stage chronic kidney disease. We did not exclude this cat from further analysis because the disease course was early, our sample size was small, and preliminary beta diversity analyses did not identify this cat as being an outlier amongst the other samples. All baseline diagnostics were performed at Oregon Veterinary Diagnostic Laboratory at Oregon State University Carlson College of Veterinary Medicine or submitted to IDEXX Laboratories Inc. (Westbrook, Maine, USA). The Feline Respiratory Panel performed at the University of California Davis School of Veterinary Medicine Real-time PCR Research and Diagnostics Core Facility was used to screen all cats for primary bacterial and viral FURTD pathogens.

### Sample collection and processing of gut and nasal microbiomes

Cats were sampled between January and March 2019. Cohabiting cats were examined on the same day and in their residence to minimize stress induced responses. Separate conjunctival, nasal, and oropharyngeal swabs were collected using sterile synthetic swabs with plastic handles (Copan®, FLOQSwabsTM, Brescia, Italy), and were pooled in 5mL of sterile phosphate-buffered saline (PBS) and stored at -80°C until extraction (7,30,33). From each cat, 4 mL of blood was collected via peripheral venipuncture for chemistry panel (serum), complete blood count (whole blood in EDTA), and FeLV/FIV/HW testing (serum). Fecal samples were collected using a fecal loop for microbiome sequencing, intestinal parasite screening (i.e. sugar flotation), and respiratory parasite screening (i.e. lung worm) (34). All samples were stored at 4°C until processing within 24 hours of collection. Blood and feces were transferred to the diagnostic laboratory for analyses. Swabs were vortexed vigorously and 2mL aliquots of PBS were removed and stored separately. Both aliquots were centrifuged at 2500g for 20 minutes (30). The resulting pellets were resuspended in 500μL PBS for DNA extraction, and stored at -80°C until extraction. Original swabs and remaining PBS were submitted for primary FURTD pathogen testing.

### 16S DNA extraction and processing

All samples were processed on the same day. At the time of DNA extraction, the swabs in PBS were removed from the freezer, thawed at 37°C, and vortexed vigorously for 15 seconds. The swab was removed, and the remaining PBS was centrifuged at 2500g for 20 minutes. The resulting pellet was resuspended in ATL buffer, and DNA was extracted using the Qiagen Inc. (Hilden, Germany) DNeasy PowerSoil Kit. The amplicons were cleaned, indexed, and sequenced using the Illumina MiSeq platform. “Universal” 16S sequencing primers were utilized from the Earth Microbiome Project 16S Illumina Amplicon Protocol (35). Forward (5’-GTGYCAGCMGCCGCGGTAA-3’) (36) and reverse primers (5’-GGACTACNVGGGTWTCTAAT-3’) (37) amplified the V4 hypervariable region.

### Amplicon Sequence Variant (ASV) determination and rarefication

A total of 15 gut samples and 17 combined nasopharyngeal samples were provided for sequencing. ASVs were determined using the <u>D</u>ivisive <u>A</u>mplicon <u>D</u>enoising <u>A</u>lgorithm (DADA2; *version 1.16.1*) pipeline (38) in R (*Joy In Playing*; version 3.5.0). The forward primer and reverse-reverse complement primer were trimmed from the forward reads, and the reverse and forward-reverse complement were trimmed from the reverse reads. Forward reads were trimmed at 250 basepairs and reverse reads were trimmed at 200 basepairs after examining quality scores. Convergent error model parameterization and sample inference was estimated for forward and reverse reads (38). Paired ends were merged, and chimeras were excluded (38). Any sequence that was not taxonomically annotated as Bacteria or Archaea was excluded from further analyses. Collector’s curves were used to determine appropriate rarefaction depth. Because the number of unique ASVs per sample were asymptotic at the minimum read depth, nasal and gut samples were rarified without replacement (*set.seed*(*77*)) to the sample with the lowest sequencing depth (41,081 and 36,494 reads respectively). Rarefaction resulted in final ASV tables with 949 unique gut ASVs and 989 unique nasopharyngeal ASVs.

### Phylogenetic tree construction and determination of Cladal Taxonomic Units (CTUs)

Analysis was conducted separately for gut and nasal communities. ASVs were combined with full length reference 16S rDNA sequences from the All Species Living Tree Project (SILVA version 1.32) (39). ASVs and reference sequences were aligned using mothur (40) (version 1.45.0). ASVs which did not align were disregarded from further analysis. A generalizable time-reversible phylogenetic model was constructed from the combined SILVA full length and ASV sequences using FastTree *(-gtr -nt*) (version 2.1.10). The resultant tree was midpoint rooted using the phangorn R package, and the reference sequences were subsequently pruned from the resultant phylogenetic tree. CTUs were determined by using the Cladal Taxonomic Unit (ClaaTU) Algorithm as described previously (31). In brief, ClaaTU conducts a root-to-tip traversal of the phylogenetic tree and quantifies the abundance of every monophyletic lineage within the reference tree within each sample. Taxonomy of each clade was assigned by determining the most specific Linnaean taxonomic label that was shared by all cladal descendants.

### Phylogenetic clustering of microbial features associated with clinical signs

To determine microbial features (i.e. ASVs or CTUs) which were predicted by the presence or absence of clinical signs, we modeled microbial features as a function of host status using the negative binomial distribution with the *nb.glm()* function. If the model failed to converge, we did not include it in the final results. We considered only microbial features which occurred in at least >40% of individuals for models. To correct for multivariate hypothesis testing, a false discovery rate correction (*q<0.05*) was applied. To determine presence of phylogenetic clustering, we calculated pairwise phylogenetic distance between all significant microbial features that were associated with clinical signs. We then calculated a random pairwise phylogenetic distance between pairs of random nodes on the phylogenetic tree. To compare distributions, we used the Kolmogorov Smirnov Test. We tested for differences in average distance between features associated with clinical signs vs. random microbial features using a Wilcox Ranked Sum Test. Features were visualized on the phylogenetic tree using the package ggtree (version 3.1.2.992).

## Results

### Inflammatory markers differ in cats with and without clinical signs of chronic FURTD

Cats with chronic FURTD signs were recruited based on a clinical history of the presence of chronic (> 1 year duration) signs (n = 11) from 8 different households (**Table 1**). When available, cohabitating housemates which were apparently normal (n = 6) were also included, resulting in a total of 17 individuals included in the final study. Cats ranged in age from 4 - 14 with a mean age of 7.4 years. There were 10 male neutered and 7 female spayed cats. The most common clinical signs of cats in the FURTD group reported by owners were nasal discharge (7/11), sneezing (7/11), ocular discharge (6/11), and ocular crusts (5/8).

We hypothesized that markers of inflammation would differ in cats with and without clinical signs of FURTD due to systemic inflammatory effects of disease (40). Although cell counts were within normal reference ranges for all cats, the number of neutrophils / uL of whole blood was significantly increased in the FURTD group compared to controls (**Figure 1A**), consistent with an inflammatory response in the FURTD group (Wilcoxon Rank Sum Test; *p = 0.024*). We also found that albumin levels were significantly lower in FURTD cats compared to controls (*p = 0.029*), consistent with the expected negative protein acute phase protein response (**Figure 1B**). No significant difference was found between cases and controls for other blood values commonly associated with the acute inflammatory response such as differences in globulins, banded neutrophils, and other leukocytes.

**Figure 1:**
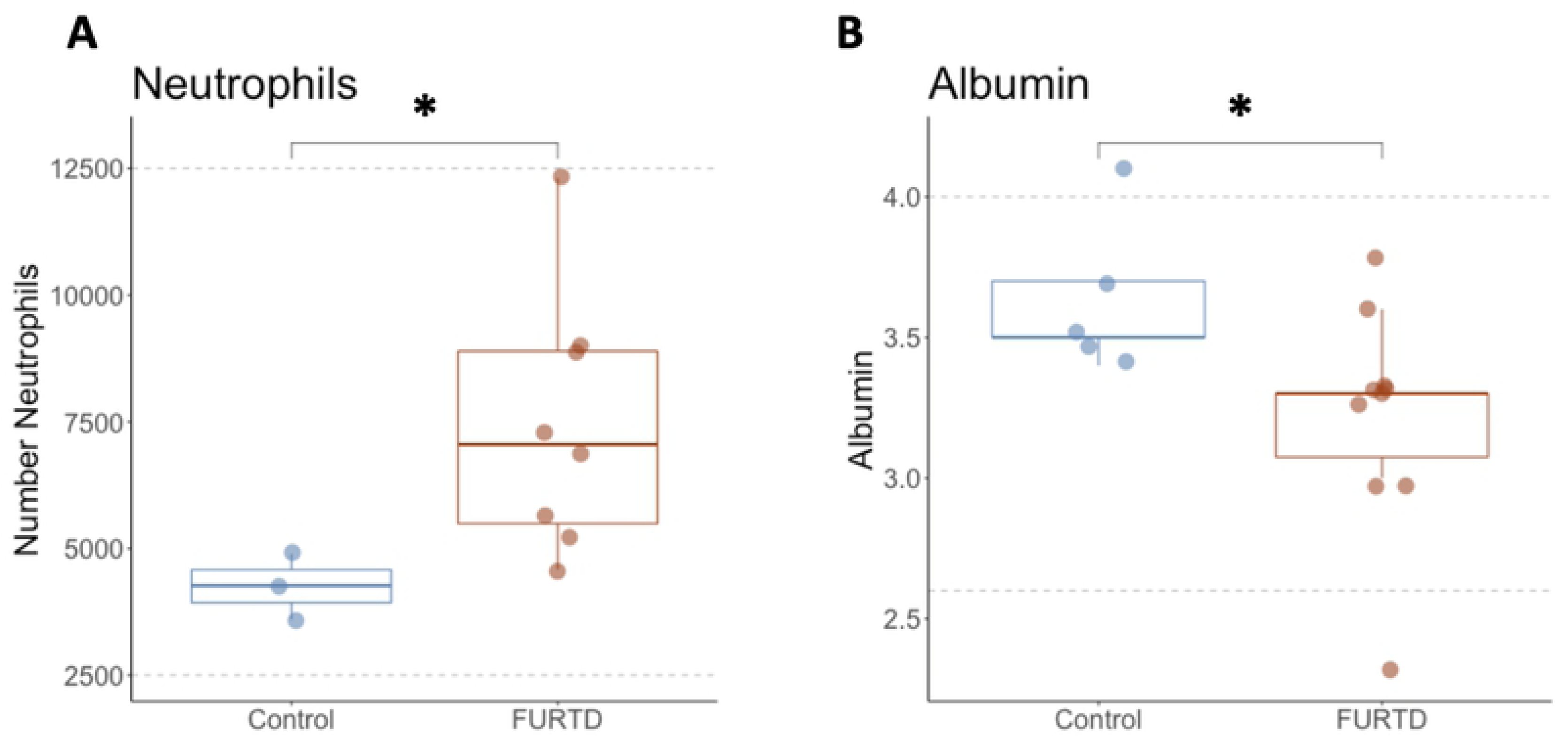
Markers of inflammation correlate with FURTD status. Absolute number of neutrophils (1A) and albumin (1B) levels are displayed for cases and controls. Box plots summarize the median, lower quartile and upper quartile for each value. Laboratory reference ranges for neutrophil numbers and albumin levels are bounded by grey dashed lines.

### Gut and nasal microbiome features significantly associate with presence of clinical signs

Next, we wished to explore if there were gut and nasal microbiome features (i.e. ASVs or microbial lineages) that associate with the presence of clinical signs. Using a recently developed ecophylogenetic framework (31, 41), we modeled each monophyletic lineage, or <u>c</u>ladal taxonomic <u>u</u>nit (CTU) as function of host status to identify those lineages of microbes significantly associated with chronic clinical signs. CTUs are taxonomic label independent, thus allowing for association of phylogenetic groups to host health status at any phylogenetic level. We also modeled each ASV as a function of host status to identify putative biomarker ASVs that might be used as indicators of host health status.

Overall, there were a total of 1,897 gut microbial features (ASVs and CTUs), of which 123 (6%) were significantly associated with the presence of chronic FURTD signs after correction for multiple hypothesis testing (*fdr < 0.05*) (**Figure 2**). Taxonomic labels most commonly associating with control individuals included 5 lineages of Porphyromonadaceae, 4 lineages of Lachnospiraceae, Prevotellaceae, and Ruminococcaceae. Microbial families most commonly associating with cats with chronic FURTD signs included 13 different lineages of Bacteroidaceae, 12 lineages of Clostridiales Family XI, Lachnospiraceae, and Burkholderiaceae.

**Figure 2:**
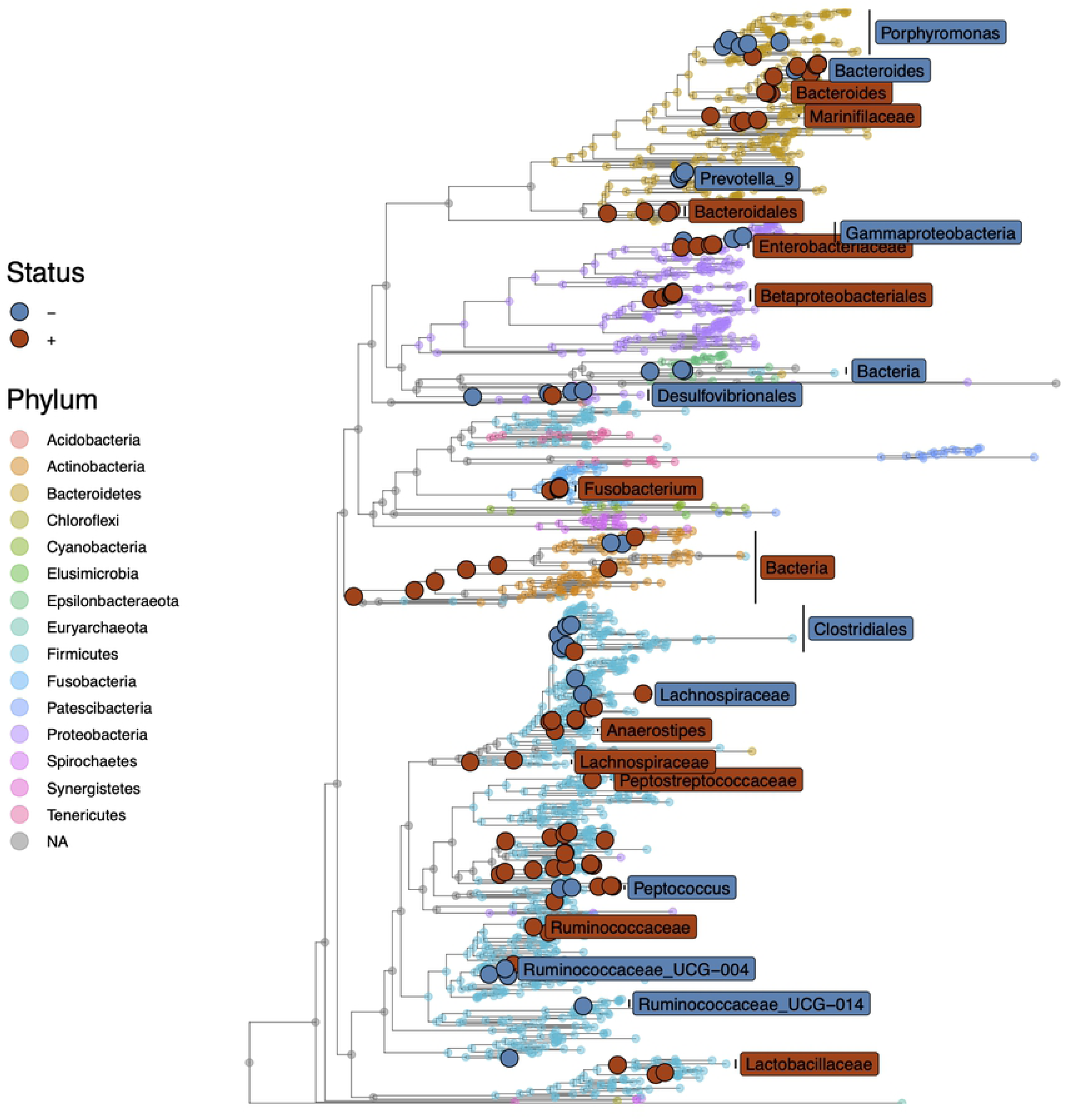
Gut microbiota features significantly associate with the presence of clinical signs. Each microbial feature was modeled as a function of nasal or ocular clinical signs. After correction for multiple hypothesis testing (*fdr* < 0.05), we visualized phylogenetic distribution of significant associations on the gut microbial tree. Labels are colored to indicate their association with the presence (*red*) or absence (*blue*) of chronic FURTD clinical signs. The bacterial taxonomic phylum label is indicated by tree tip colors.

Of the total 1,977 nasal microbiome features (i.e. CTUs and ASVs), only 75 (3%) were significantly associated with the presence of chronic FURTD signs (*fdr < 0.05*) (**Figure 3**). The most common microbial features which significantly associated with control individuals included Porphyromonadaceae (*n = 6*), Peptostreptococcaceae (*n = 4*), Pasteurellaceae (*n = 4*), Prevotellaceae (*n = 3*), Clostridiales Family XII (*n = 3*), and Spirachaetaceae (*n = 3*). Lineages most commonly associated with cases were included the taxonomic labels Actinomycetaceae (*n = 4*), Porphyromonadaceae (*n = 4*), Pasteurellaceae (*n = 4*), and Moraxellaceae (*n = 4*).

**Figure 3:**
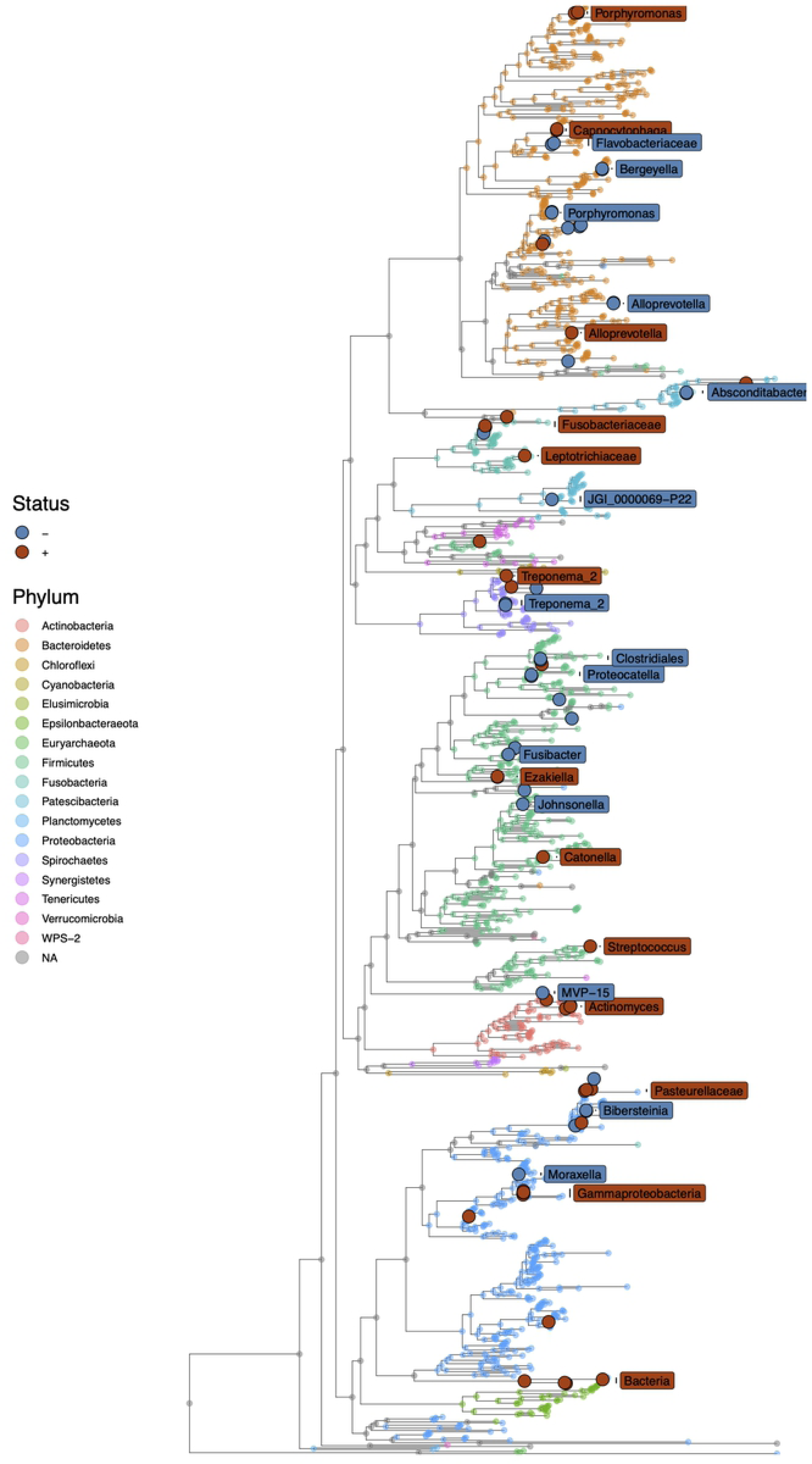
Nasal microbiota features significantly associate with the presence of nasal and ocular clinical signs. Each microbial feature was modeled as a function of clinical sign presence or absence. After correction for multiple hypothesis testing (*fdr* < 0.05), we visualized phylogenetic distribution of significant associations on the nasal microbial tree. Labels are colored to indicate their association with clinical signs. The bacterial taxonomic phylum label is indicated by tree tip colors.

### Gut microbiome lineages show conserved patterns of association with host disease status

Of the 123 microbial features that had significant associations with the host, there were 37 distinct monophyletic microbial lineages which showed patterns of association within the gut microbiome. We hypothesized that descendants within these microbial lineages may show similar association patterns with host status. We reasoned that related groups of microbes are more likely to have more related functional capacity, and thus are more likely to share similar patterns of association with the host. Because of this, we hypothesized that there would be positions on the phylogenetic tree that showed phylogenetic clustering (i.e. “hot spots” of association) with the host status.

We found that microbial features of the gut significantly associating with clinical signs exhibited evidence of phylogenetic clustering. Microbial lineages and ASVs positively and negatively associating with clinical signs have a distinct distribution compared to random as well as one another (KS Test; *p << 0.01*). The mean pairwise phylogenetic distance between two random nodes on the gut phylogenetic tree was significantly greater (*dist_μ_rand_ =1.18; SD_rand_ = 0.42*) than distance between nodes that either positively (*dist_μ+_ = 1.0; SD_+_= 0.43*) or negatively (*dist_μ-_ = 1.04; SD_μ-_ = 0.43*) associated with clinical signs (Wilcoxon rank sum test *p<<0.01*). Phylogenetic clustering is consistent with a conserved phylogenetic core of clades that are present in cats with and without clinical signs compared to random. The pairwise average distance between microbial features which positively associated with clinical signs was significantly less than pairwise distances between clades which negatively associated with clinical signs (**Figure 4**) (*p<<0.01*), and also had an additional peak of short pairwise distances. This could result from population expansion of a small set of closely related microbes during disease.

**Figure 4:**
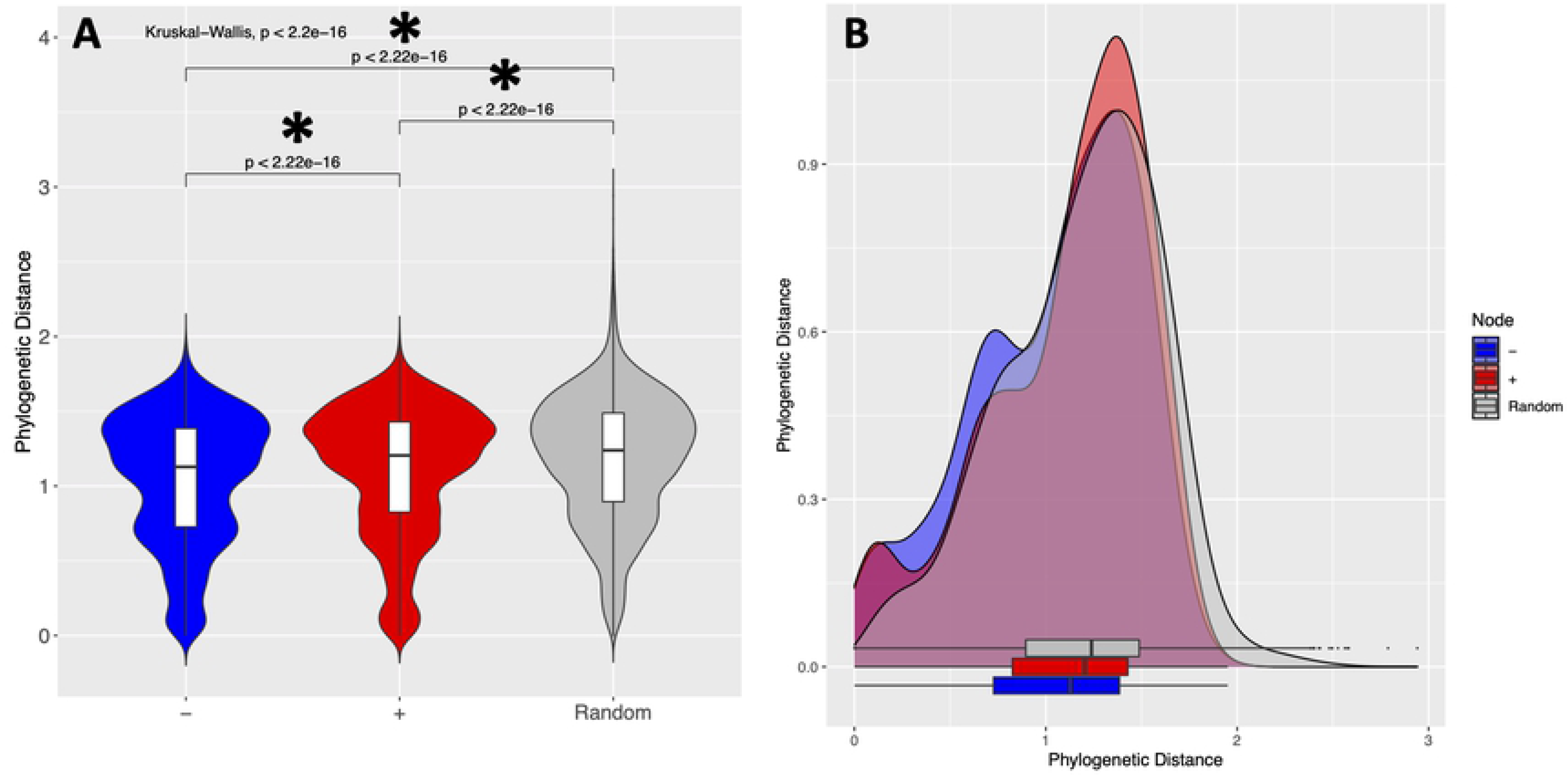
Gut microbiome features associating with clinical signs shows evidence of phylogenetic clustering. The pairwise phylogenetic distance between all pairs of microbial features positively (*red*) or negatively (*blue*) associating with clinical signs was calculated and compared to a random sampling of pairwise distances (*grey*). **4A**: Violin plots showing pairwise phylogenetic distances between clades significantly associating with clinical signs. Significant differences between groups were tested using the Wilcoxon rank sum test and p-values are displayed above bars. **4B**: Density plots of pairwise distances between significant microbial features compared to random.

Examining the association of each descendant within lineages significantly associated with host status, we found some striking examples of conservation of host microbe associations (**Figure 5**). For example, several related ASVs taxonomically annotated as Porphyromonas (Phyla Bacteroidetes; Order Bacteroidales) are all increased in control cats compared to FURTD individuals. Conversely, we found that in cats with clinical signs, there is a relative increased abundance of a particular clade taxonomically annotated as the family Lactobacillaceae. This lineage contained 9 different members, 8/9 of which were annotated as *Lactobacillus*, and 1/9 of which was annotated as *Pediococcus*.

**Figure 5:**
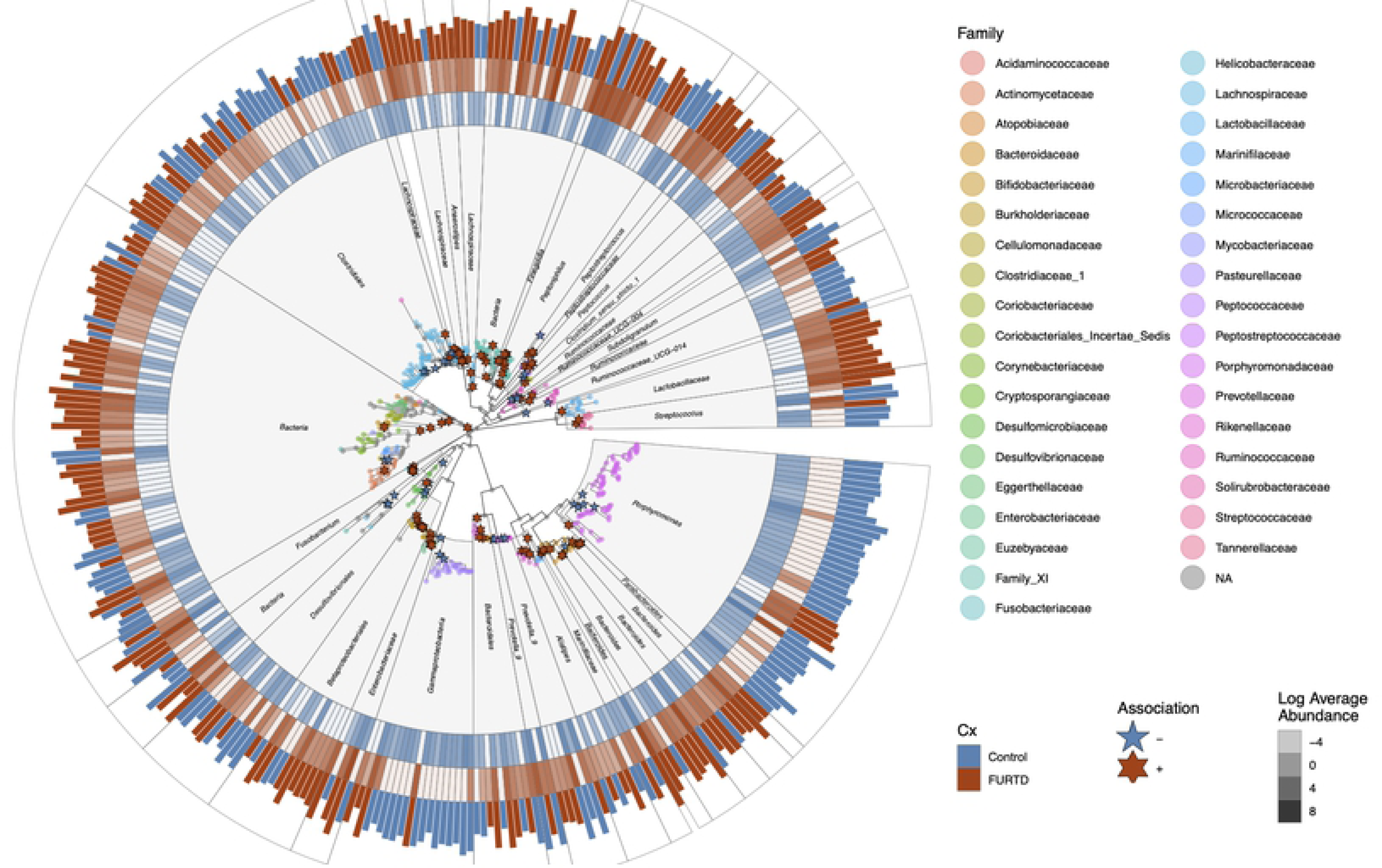
Monophyletic lineages within the gut microbiome reveal distinct cladal patterns of association with clinical signs. Monophyletic lineages significantly associating with clinical signs were subset and displayed. Microbial taxonomic family is indicated by node colors. Significant associations with FURTD or control individuals indicated with red and blue stars respectively. Tiles represent the log transformed average abundances of each ASV in individuals without (*blue*) and with (*rust*) clinical signs. Bars along the outside of the figure represent the highest average abundance of controls compared to FURTD individuals. Bars are color coded by which group showed higher abundances.

### Nasal microbiome lineages show conserved patterns of association with host disease status

Of the 75 microbial features of the nasal microbiome that had significant associations with chronic FURTD signs, there were only a total of 27 distinct microbial lineages that were associated with host disease status. As before, we hypothesized that descendants within these microbial lineages may show similar association patterns with host status. We reasoned that related groups of microbes are more likely to have more related functional capacity, and thus are more likely to share similar patterns of association with the host. Because of this, we hypothesized that there would be positions on the phylogenetic tree that showed phylogenetic clustering (i.e. “hot spots” of association) with the host status In contrast to the gut microbiome, microbial features of the nasal microbiome significantly associating with clinical signs do not exhibit evidence of phylogenetic clustering. Mean pairwise phylogenetic distance between two random nodes on the nasal phylogenetic tree was less (dist*_μ_rand_* = 1.31; SD*_rand_* = 0.42) than nodes positively (dist*_μ+_* = 1.38; SD*_+_* = 0.45) and negatively (dist*_μ-_* = 1.34; SD*_μ-_* = 0.42) associated with clinical signs (Wilcoxon rank sum test *p<0.01*) (**Figure 6**). The lack of evidence for phylogenetic clustering is consistent with increased variability present in nasal microbiome communities. This result is an interesting contrast from the gut microbiome and may suggest that nasal microbes that link to presence of active clinical signs are phylogenetically dispersed or less subject to vertical evolution as compared to those traits that link nasal microbial lineages to clinical signs.

**Figure 6:**
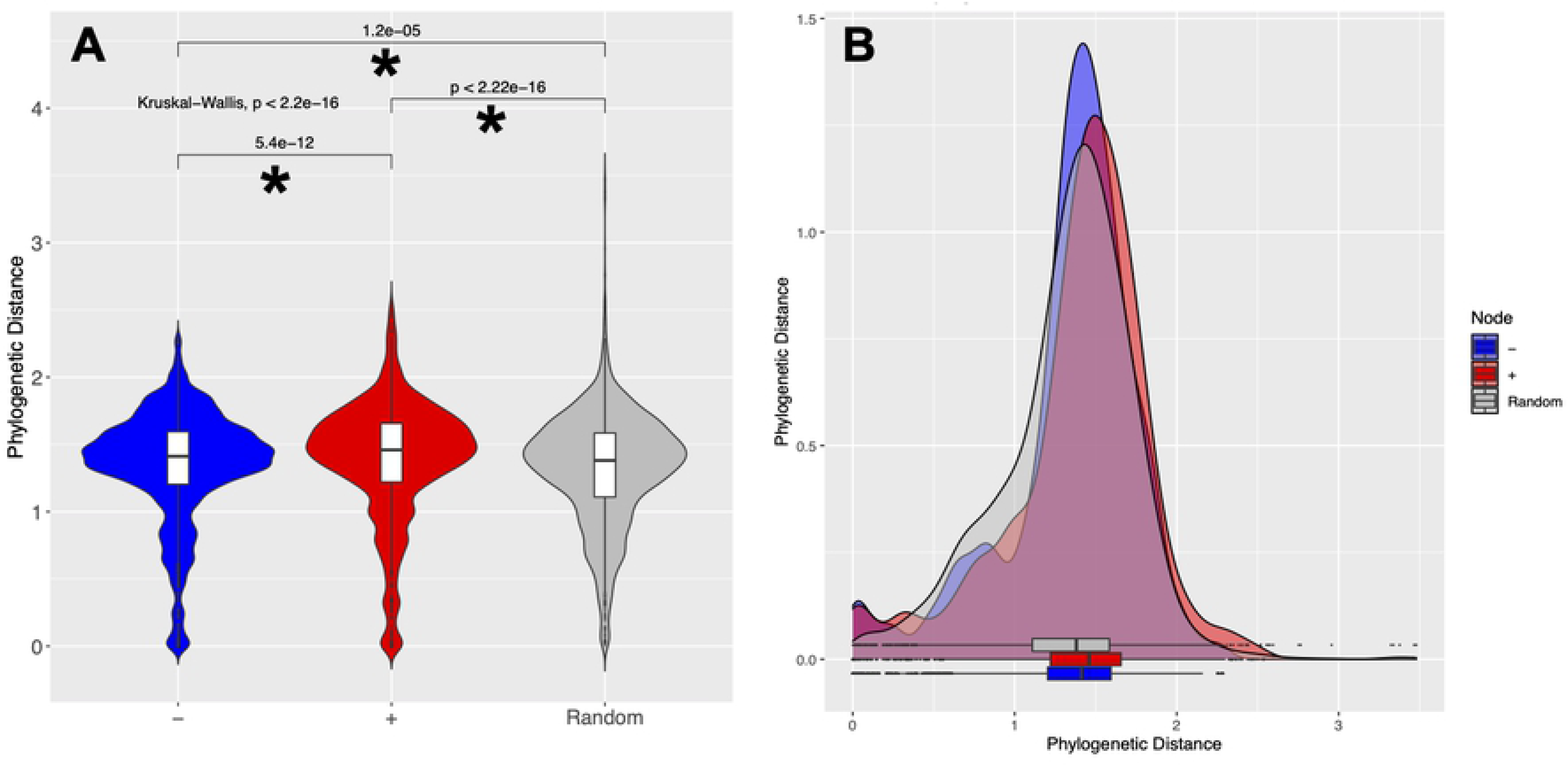
Nasal microbiome features positively associating with clinical signs do not show evidence of phylogenetic clustering. The pairwise phylogenetic distance between all pairs of microbial features positively (*red*) or negatively (*blue*) associating with clinical signs was calculated and compared to a random sampling of pairwise distances (*grey*). **6A:** Violin plots showing pairwise phylogenetic distances between clades significantly associating with clinical signs. Significant differences between groups were tested using the Wilcoxon rank sum test and p-values are displayed above bars. **6B:** Density plots of pairwise distances between significant microbial features compared to random.

Considering cladal information reveals unique insight into patterns associating with nasal microbiome features which associate with clinical signs. For example, consider the heterogeneity of association of clades with the taxonomic label within the Class Gammaproteobacteria and Family Moraxellaceae (**Figure 7**). One lineage taxonomically annotated as Moraxellaceae has descending lineages which all show increased abundances in cats with absent clinical signs, while another clade within Gammaproteobacteria contains six descending lineages of *Moraxella* displaying association with presence of clinical signs.

**Figure 7:**
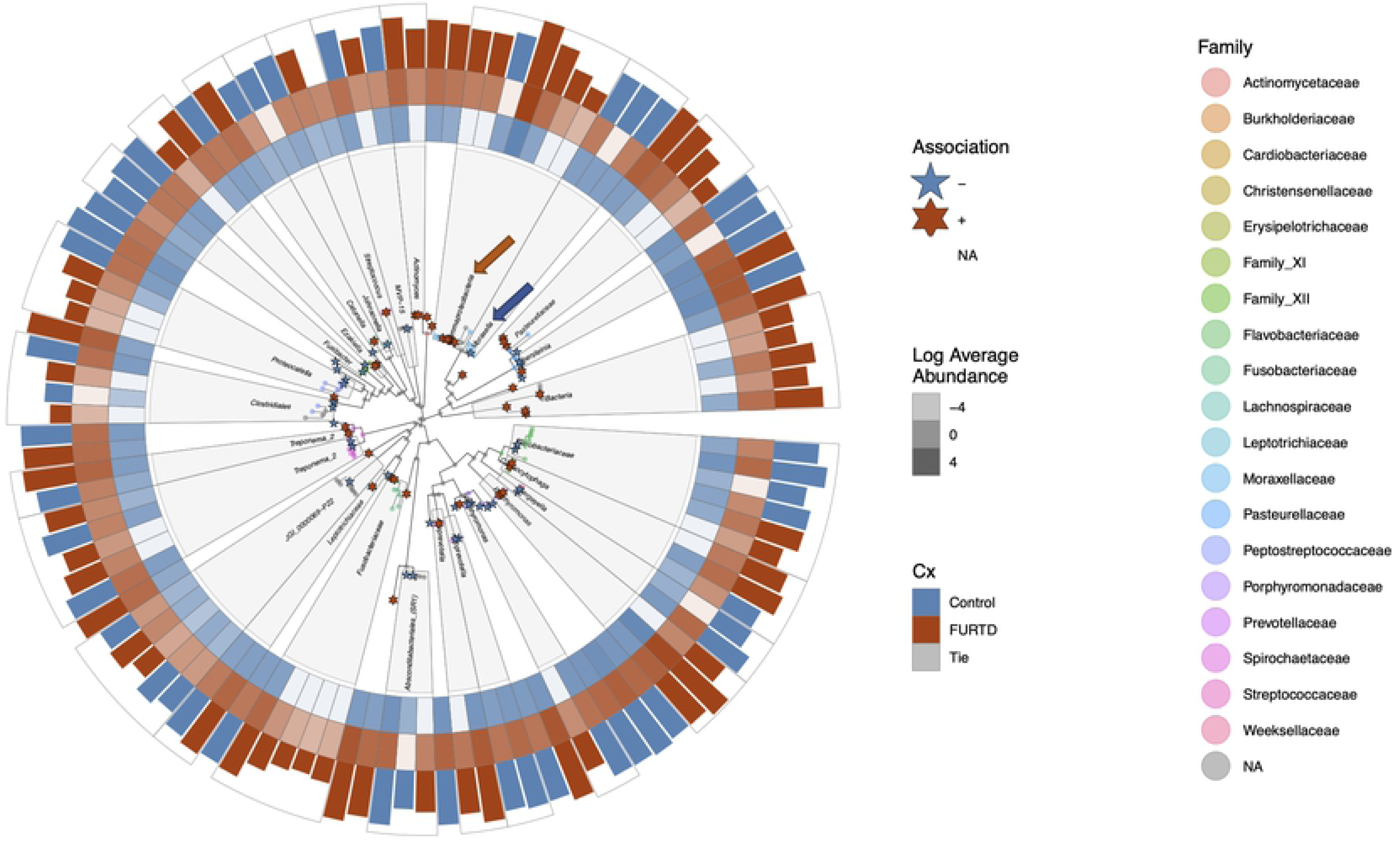
Monophyletic lineages within the nasal microbiome reveal distinct cladal patterns of association with clinical signs. Monophyletic lineages significantly associating with clinical signs were subset and visualized separately. Microbial taxonomic family is indicated by node colors. Significant associations with clinical signs are indicated with stars. Tiles represent the log transformed average abundances of each ASV in individuals without (*blue*) and with (*rust*) clinical signs. One lineage taxonomically annotated as Moraxellaceae has descending lineages which all show increased abundances in cats with absent clinical signs (blue arrow), while another clade within Gammaproteobacteria contains six descending lineages of *Moraxella* displaying association with presence of clinical signs (red arrow).

## Discussion

Understanding the differences in community compositions between cats with and without chronic signs of FURTD may provide new insight into how the microbial community interacts with the primary pathogens leading to development of chronic clinical disease. We identified specific microbial features that linked with presence or absence of chronic clinical signs. Understanding the taxonomic community composition and functional capacity of the feline upper respiratory tract and gut microbiome and their respective roles in chronic FURTD pathogenesis may provide new lines of study that could produce novel therapeutics for cats suffering from chronic FURTD signs. For example, if the gut microbiome is a mediator of clinical signs, understanding which microbes may mitigate viral effects may lead to new probiotics that may lessen chronic clinical signs of infectious FURTD. In addition, we identified several microbial features that may hold high information content at being able to predict the presence of chronic clinical signs in cats. Future work could determine the prognostic potential of the likelihood that cats would develop chronic clinical course upon initial infection given the presence or absence of these “biomarker clades” and ASVs.

An overall inflammatory state provides a putative mechanism for restructuring gut and nasal microbial communities. As expected, we found that cats with chronic FURTD signs had increased levels of inflammation as indicated by significantly higher neutrophil counts and reduction in albumin, a negative acute phase protein. It is probable that during an inflammatory state, the host tends to filter for particular monophyletic lineages of microbes within the gut and nasal passages via conserved microbial traits. Our study also suggests that clinically relevant changes in absolute neutrophil number and albumin still exist within normally established reference ranges.

Our findings link the gut microbiome to the presence of chronic FURTD signs. Microfold cells (M-cells) are cells that sample microbes within the gut and lay within the epithelia overlying the gut-associated lymphoid tissues (GALT) (42). M-cells allow for transcytosis of luminal antigens to mononuclear phagocytes, thereby inducing mucosal immune responses such as IgA production (42). Recently, cells with the typical features of M-cells have been found within murine nasal passage epitheliums, overlying a novel Nasal Associated Lymphoid Tissue (NALT). NALT enables independent sampling of pathogens within the respiratory tract (42). It is well known that there is crosstalk between different mucosal systems within the body, so IgA production in the NALT may affect gut microbiomes and vice versa (43). For example, many studies have shown cross mucosal interaction at distant sites, such as mucosal vaccination leading to protection at another mucosal surface, infection with a virus at one mucosal site resulting in IgA secretions at distal sites, and greater risk of respiratory allergies in neonates that are put on a course of antibiotics (43).

We discovered several microbial clades within the gut which are known to have probiotic properties mediating the gut - respiratory axis in higher abundances in cats with chronic signs compared to those without. This apparent paradox has been described previously in studies that monitor enteric gut microbiome population response upon infection with respiratory virus (44). For example, we discovered a higher number of microbes taxonomically annotated as *Lactobacillus* in cats with signs of chronic FURTD. *Lactobacillus* has classically been associated with protection from respiratory viral infection in several disease models due to its established role as a probiotic mediator of the gut-lung axis. Changes within enteric microbial populations upon infection with respiratory virus is less studied, but prior studies speculate that host recruitment of microbes with anti-inflammatory properties may provide a host resilience to viral infection. For example, expansions of murine enteric populations of *Lactobacillus* have been shown to increase upon infection with influenza virus (44), and decreasing *Lactobacillus* numbers upon respiratory viral infection has been shown to increase risk of death in a mouse model (45). The authors speculate that increases in gut *Lactobacillus* numbers may therefore be a host adaptive host response to viral infection. Administration of *Lactobacillus* strains in mice can decrease asthmatic symptoms (46–49), decrease airway inflammation and alveolar damage (49, 50), and provide protection against respiratory viral infection (23,45,51). Thus, we might expect *Lactobacillus* numbers to increase in those animals with chronic FURTD signs compared to controls as an adaptive response.

We found a relatively higher abundance of microbes within Porphyromonas within the order Bacteroidales in control individuals. Reduction in relative abundance of Bacteroidales has been described with respiratory viral infection in mouse models previously. For example, there was a dramatic reduction of gut Bacteroidales numbers upon infection with influenza A virus (44), but the authors did not go beyond taxonomic labels to further describe if all members of Bacteroidales taxa exhibited similar reductions, or if some taxonomic labels exhibited opposite effects. In our study, reduction of Bacteroidales was driven largely by a diffuse association with subtending lineages of the Porphyromonas. Other members within the Bacteroidales family did not manifest similar patterns of association with clinical signs.

We found several lineages within the nasal microbiome that were linked to the presence of chronic FURTD course. One such lineage within labeled as the class Gammaproteobacteria had opposite associations with host status – one group of descending lineages (labeled as *Moraxella*) had negative associations with host disease status, while other descending lineages (also labeled as *Moraxella*) within the same class positively associated with clinical signs. *Moraxella* isolates have been noted for their heterogeneity in ability to cause clinical signs (52). While some lineages of *Moraxella* are commensals, others are considered important respiratory pathogens that can worsen other diseases of the respiratory tract such as chronic obstructive pulmonary disease (53). In humans, recent acquisition of one of two distinct lineages of *Moraxella catarrhalis* has been linked to exacerbations of airway inflammation and increased clinical signs (53–55). Some *Moraxella* strains in humans produce β-lactamase, and so are resistant to ampicillin. In addition, *Moraxella* resists antibiotic treatment by its ability to form biofilms, recently implicated in recurrent and chronic otitis media and upper respiratory signs in children (56). Another clade which is strongly linked to presence of FURTD clinical signs is *Fusobacterium*, which is also one of the most common isolates in humans with chronic sinusitis (57).

Our ecophylogenetic approach yields unique discovery that examination of taxonomic or ASVs alone cannot. Considering each and every clade’s association with host status can reveal divergent functional groups. For example, consider if the clade *Moraxella* had been collapsed at genus level to a single label and then assessed for association with host status. Because some descendants have a positive association with disease and others have a negative association with disease, if we were to consider the group as a whole, it is possible that neither would have been significantly associated with host status. Also, an ecophylogenetic approach yields groups of descendants that likely have conserved traits of association with host disease status that could be followed up upon with comparative genomics approaches to identify specific bacterial mechanisms of interaction with host.

Our work suffers from several limitations. First, while our study benefits from being able to rapidly determine microbial features that associate with FURTD clinical signs, it is limited by its cross sectional nature. Because of this, the directionality of cause and effect cannot be determined. It is possible that the microbes found in higher abundances in FURTD cats contribute to severity of clinical signs. Conversely, it is possible that clinical signs leads to selection of different microbial communities. Second, we are limited by the number of cats in our study – future work should seek to recruit larger cohorts in order to identify the robustness of associations determined here. Third, our work is limited by the lack of definitive diagnosis for the cats in question. It is possible that there were other causes besides infectious FURTD leading to chronic nasal signs on our recruited population. Additional studies should seek to definitively rule out the presence of other diseases leading to chronic upper respiratory tract signs through use of additional diagnostics (e.g. biopsy and rhinoscopy). Unfortunately, due to cost constrains and invasive nature of rhinoscopy, use of these tests were not feasible in the current study. Last, we only considered the bacterial component of the nasal and gut microbiomes in regards to FURTD. Future work should seek to characterize fungal communities within nasal passages given that fungal organisms such as *Cryptococcus* can be a cause of infectious FURTD. It is also possible that other pathogens contribute to the disease process that have yet to be described. Future work should consider virome as well as microbiome sequencing to identify other possibilities of host viral interactions that lead to long term disease sequalae.

The gut and nasal microbiome may be a useful prognostic indicator for which cats may develop chronic clinical disease upon acute infection. Future work should seek to determine the reason some cats develop chronic clinical disease and some do not. For example, do the microbial lineages which were increased in control compared to FURTD cats have “protective” effects against development of chronic clinical course? Are microbes that are associated with chronic disease present upon initial infection with the primary pathogen, or do they establish after acute infection of the primary pathogen? We were not able to answer these questions because our work only considered one time point, but future studies which establish how microbial communities respond to initial infection and then develop during chronic clinical disease course will shed light on cause and effect. Future work describing the interactions between the gut and respiratory microbiomes could provide the basis for novel therapeutics. Ultimately, protective probiotic strains given to shelter animals on intake may reduce the amount of morbidity associated with initial infection, as well as to decrease the rate of chronic clinical disease.

## Acknowledgments

We thank Lucie Crane for helpful discussions on overall paper narrative. Thank you Kristin Kasschau for help with DNA extractions. Thank you Sam Lastufka for help with sample collection and data entry. Thank you Martha F. Witkop for providing comments. The Morris Animal Foundation (Grant D19FE-808) to BRB and RH, the National Science Foundation (Grant 1557192), and institutional funds to TJS supported this work.

## Author contributions

HKA processed raw sequencing data, performed microbial statistical tests, produced data visualization results, and wrote the main manuscript text. BRB, SMD, and RH collected samples. BRB and RH did all veterinary examination procedures. BRB and SMD performed laboratory work. HKA and SMD entered data. BRB and RH conceptualized the project. SML performed statistical tests comparing blood work values. BRB, RH, and TJS oversaw the analysis and writing process. All authors contributed to revisions and approved the final version.

## Data Availability

Raw sequences used to generate these results will be available on the National Center for Biotechnology Information’s Sequence Read Archive under project identification number PRJNA835156 upon publication.

## Notes

### Competing Interest Statement

The authors have declared no competing interest.

## References

1. Burns RE, Wagner DC, Leutenegger CM, Pesavento PA. Histologic and molecular correlation in shelter cats with acute upper respiratory infection. Journal of clinical microbiology. 2011;49(7):2454–60.

2. Dinnage JD, Scarlett JM, Richards JR. Descriptive epidemiology of feline upper respiratory tract disease in an animal shelter. Journal of feline medicine and surgery. 2009;11(10):816– 25.

3. Lappin M, Blondeau J, Boothe D, Breitschwerdt E, Guardabassi L, Lloyd D, et al. Antimicrobial use guidelines for treatment of respiratory tract disease in dogs and cats: antimicrobial guidelines working group of the International Society for Companion Animal Infectious Diseases. Journal of veterinary internal medicine. 2017;31(2):279–94.

4. Quimby J, Lappin M. Feline focus: Update on feline upper respiratory diseases: introduction and diagnostics. Compendium (Yardley, PA). 2009;31(12):E1–7.

5. Cohn LA. Feline respiratory disease complex. Veterinary Clinics: Small Animal Practice. 2011;41(6):1273–89.

6. Bannasch MJ, Foley JE. Epidemiologic evaluation of multiple respiratory pathogens in cats in animal shelters. Journal of Feline Medicine & Surgery. 2005;7(2):109–19.

7. Dorn ES, Tress B, Suchodolski JS, Nisar T, Ravindran P, Weber K, et al. Bacterial microbiome in the nose of healthy cats and in cats with nasal disease. PloS one. 2017;12(6):e0180299.

8. Grahn BH, Sisler S, Storey E. Qualitative tear film and conjunctival goblet cell assessment of cats with corneal sequestra. Veterinary ophthalmology. 2005;8(3):167–70.

9. Burgesser KM, Hotaling S, Schiebel A, Ashbaugh SE, Roberts SM, Collins JK. Comparison of PCR, virus isolation, and indirect fluorescent antibody staining in the detection of naturally occurring feline herpesvirus infections. Journal of Veterinary Diagnostic Investigation. 1999;11(2):122–6.

10. Hara M, Fukuyama M, Suzuki Y, Kisikawa S, Ikeda T, Kiuchi A, et al. Detection of feline herpesvirus 1 DNA by the nested polymerase chain reaction. Veterinary microbiology. 1996;48(3–4):345–52.

11. Lim CC, Cullen CL. Schirmer tear test values and tear film break-up times in cats with conjunctivitis. Veterinary ophthalmology. 2005;8(5):305–10.

12. Low HC, Powell CC, Veir JK, Hawley JR, Lappin MR. Prevalence of feline herpesvirus 1, Chlamydophila felis, and Mycoplasma spp DNA in conjunctival cells collected from cats with and without conjunctivitis. American journal of veterinary research. 2007;68(6):643– 8.

13. Rampazzo A, Appino S, Pregel P, Tarducci A, Zini E, Biolatti B. Prevalence of Chlamydophila felis and feline herpesvirus 1 in cats with conjunctivitis in northern Italy. Journal of veterinary internal medicine. 2003;17(6):799–807.

14. Volopich S, Benetka V, Schwendenwein I, Möstl K, Sommerfeld-Stur I, Nell B. Cytologic findings, and feline herpesvirus DNA and Chlamydophila felis antigen detection rates in normal cats and cats with conjunctival and corneal lesions. Veterinary ophthalmology. 2005;8(1):25–32.

15. Cullen CL, Wadowska DW, Singh A, Melekhovets Y. Ultrastructural findings in feline corneal sequestra. Veterinary ophthalmology. 2005;8(5):295–303.

16. von Bomhard W, Polkinghorne A, Huat Lu Z, Vaughan L, Vögtlin A, Zimmermann DR, et al. Detection of novel chlamydiae in cats with ocular disease. American journal of veterinary research. 2003;64(11):1421–8.

17. Sandmeyer LS, Waldner CL, Bauer BS, Wen X, Bienzle D. Comparison of polymerase chain reaction tests for diagnosis of feline herpesvirus, Chlamydophila felis, and Mycoplasma spp. infection in cats with ocular disease in Canada. The Canadian veterinary journal. 2010;51(6):629.

18. Townsend WM, Stiles J, Guptill-Yoran L, Krohne SG. Development of a reverse transcriptasepolymerase chain reaction assay to detect feline herpesvirus-1 latency-associated transcripts in the trigeminal ganglia and corneas of cats that did not have clinical signs of ocular disease. American journal of veterinary research. 2004;65(3):314–9.

19. Lanaspa M, Bassat Q, Medeiros MM, Muñoz-Almagro C. Respiratory microbiota and lower respiratory tract disease. Expert review of anti-infective therapy. 2017;15(7):703–11.

20. Marsland BJ, Yadava K, Nicod LP. The airway microbiome and disease. Chest. 2013;144(2):632–7.

21. Mahdavinia M, Keshavarzian A, Tobin MC, Landay A, Schleimer RP. A comprehensive review of the nasal microbiome in chronic rhinosinusitis (CRS). Clinical & Experimental Allergy. 2016;46(1):21–41.

22. Honda K, Littman DR. The microbiome in infectious disease and inflammation. Annual review of immunology. 2012;30:759–95.

23. Fujimura KE, Demoor T, Rauch M, Faruqi AA, Jang S, Johnson CC, et al. House dust exposure mediates gut microbiome Lactobacillus enrichment and airway immune defense against allergens and virus infection. Proceedings of the National Academy of Sciences. 2014;111(2):805–10.

24. Russell SL, Gold MJ, Hartmann M, Willing BP, Thorson L, Wlodarska M, et al. Early life antibiotic-driven changes in microbiota enhance susceptibility to allergic asthma. EMBO reports. 2012;13(5):440–7.

25. O’Dwyer DN, Dickson RP, Moore BB. The lung microbiome, immunity, and the pathogenesis of chronic lung disease. The journal of immunology. 2016;196(12):4839–47.

26. Ege MJ, Mayer M, Normand AC, Genuneit J, Cookson WO, Braun-Fahrländer C, et al. Exposure to environmental microorganisms and childhood asthma. New England Journal of Medicine. 2011;364(8):701–9.

27. Pakarinen J, Hyvärinen A, Salkinoja-Salonen M, Laitinen S, Nevalainen A, Mäkelä MJ, et al. Predominance of Gram-positive bacteria in house dust in the low-allergy risk Russian Karelia. Environmental microbiology. 2008;10(12):3317–25.

28. Lynch SV, Wood RA, Boushey H, Bacharier LB, Bloomberg GR, Kattan M, et al. Effects of early-life exposure to allergens and bacteria on recurrent wheeze and atopy in urban children. Journal of Allergy and Clinical Immunology. 2014;134(3):593–601.

29. Misic AM, Davis MF, Tyldsley AS, Hodkinson BP, Tolomeo P, Hu B, et al. The shared microbiota of humans and companion animals as evaluated from Staphylococcus carriage sites. Microbiome. 2015;3(1):2.

30. Vientós-Plotts AI, Ericsson AC, Rindt H, Grobman ME, Graham A, Bishop K, et al. Dynamic changes of the respiratory microbiota and its relationship to fecal and blood microbiota in healthy young cats. PloS one. 2017;12(3):e0173818.

31. Gaulke CA, Arnold HK, Humphreys IR, Kembel SW, O’dwyer JP, Sharpton TJ. Ecophylogenetics clarifies the evolutionary association between mammals and their gut microbiota. MBio. 2018;9(5):e01348–18.

32. Couch CE, Arnold HK, Crowhurst RS, Jolles AE, Sharpton TJ, Witczak MF, et al. Bighorn sheep gut microbiomes associate with genetic and spatial structure across a metapopulation. Scientific reports. 2020;10(1):1–10.

33. Hartmann AD, Hawley J, Werckenthin C, Lappin MR, Hartmann K. Detection of bacterial and viral organisms from the conjunctiva of cats with conjunctivitis and upper respiratory tract disease. Journal of feline medicine and surgery. 2010;12(10):775–82.

34. Traversa D, Di Cesare A. Diagnosis and management of lungworm infections in cats: cornerstones, dilemmas and new avenues. Journal of feline medicine and surgery. 2016;18(1):7–20.

35. Caporaso J, Ackermann G, Apprill A, Bauer M, Berg-Lyons D, Betley J, et al. EMP 16S Illumina amplicon protocol. See http://www.earthmicrobiome.org/protocols-and-standards/16s. 2018;

36. Apprill A, McNally S, Parsons R, Weber L. Minor revision to V4 region SSU rRNA 806R gene primer greatly increases detection of SAR11 bacterioplankton. Aquatic Microbial Ecology. 2015;75(2):129–37.

37. Parada AE, Needham DM, Fuhrman JA. Every base matters: assessing small subunit rRNA primers for marine microbiomes with mock communities, time series and global field samples. Environmental microbiology. 2016;18(5):1403–14.

38. Callahan BJ, McMurdie PJ, Rosen MJ, Han AW, Johnson AJA, Holmes SP. DADA2: high-resolution sample inference from Illumina amplicon data. Nature methods. 2016;13(7):581– 3.

39. Yarza P, Richter M, Peplies J, Euzeby J, Amann R, Schleifer KH, et al. The All-Species Living Tree project: a 16S rRNA-based phylogenetic tree of all sequenced type strains. Systematic and applied microbiology. 2008;31(4):241–50.

40. Schloss PD, Westcott SL, Ryabin T, Hall JR, Hartmann M, Hollister EB, et al. Introducing mothur: open-source, platform-independent, community-supported software for describing and comparing microbial communities. Applied and environmental microbiology. 2009;75(23):7537–41.

41. Sharpton TJ, Stagaman K, Sieler MJ, Arnold HK, Davis EW. Phylogenetic Integration Reveals the Zebrafish Core Microbiome and Its Sensitivity to Environmental Exposures. Toxics. 2021;9(1):10.

42. Mabbott NA, Donaldson DS, Ohno H, Williams IR, Mahajan A. Microfold (M) cells: important immunosurveillance posts in the intestinal epithelium. Mucosal immunology. 2013;6(4):666–77.

43. Tulic M, Piche T, Verhasselt V. Lung–gut cross-talk: evidence, mechanisms and implications for the mucosal inflammatory diseases. Clinical & Experimental Allergy. 2016;46(4):519–28.

44. Yildiz S, Mazel-Sanchez B, Kandasamy M, Manicassamy B, Schmolke M. Influenza A virus infection impacts systemic microbiota dynamics and causes quantitative enteric dysbiosis. Microbiome. 2018;6(1):1–17.

45. Zhang Q, Hu J, Feng JW, Hu XT, Wang T, Gong WX, et al. Influenza infection elicits an expansion of gut population of endogenous Bifidobacterium animalis which protects mice against infection. Genome biology. 2020;21(1):1–26.

46. Arrieta MC, Stiemsma LT, Dimitriu PA, Thorson L, Russell S, Yurist-Doutsch S, et al. Early infancy microbial and metabolic alterations affect risk of childhood asthma. Science translational medicine. 2015;7(307):307ra152-307ra152.

47. Kozakova H, Schwarzer M, Tuckova L, Srutkova D, Czarnowska E, Rosiak I, et al. Colonization of germ-free mice with a mixture of three lactobacillus strains enhances the integrity of gut mucosa and ameliorates allergic sensitization. Cellular & molecular immunology. 2016;13(2):251–62.

48. Wu CT, Chen PJ, Lee YT, Ko JL, Lue KH. Effects of immunomodulatory supplementation with Lactobacillus rhamnosus on airway inflammation in a mouse asthma model. Journal of Microbiology, Immunology and Infection. 2016;49(5):625–35.

49. Chunxi L, Haiyue L, Yanxia L, Jianbing P, Jin S. The gut microbiota and respiratory diseases: new evidence. Journal of Immunology Research. 2020;2020.

50. Verheijden K, van Bergenhenegouwen J, Garssen J, Bezemer G, Kraneveld A, Folkerts G. Treatment with specific prebiotics or probiotics prevents the development of lung emphysema in a mouse model of COPD. European Journal of Pharmacology. 2011;668:e12–3.

51. Zrelli S, Amairia S, Zrelli M. Respiratory syndrome coronavirus-2 response: Microbiota as lactobacilli could make the difference. Journal of Medical Virology. 2021;93(6):3288–93.

52. López-Serrano S, Galofré-Milà N, Costa-Hurtado M, Pérez-de-Rozas AM, Aragon V. Heterogeneity of Moraxella isolates found in the nasal cavities of piglets. BMC veterinary research. 2020;16(1):1–12.

53. Goldstein EJ, Murphy TF, Parameswaran GI. Moraxella catarrhalis, a human respiratory tract pathogen. Clinical Infectious Diseases. 2009;49(1):124–31.

54. Sethi S, Wrona C, Eschberger K, Lobbins P, Cai X, Murphy TF. Inflammatory profile of new bacterial strain exacerbations of chronic obstructive pulmonary disease. American journal of respiratory and critical care medicine. 2008;177(5):491–7.

55. Sethi S, Muscarella K, Evans N, Klingman KL, Grant BJ, Murphy TF. Airway inflammation and etiology of acute exacerbations of chronic bronchitis. Chest. 2000;118(6):1557–65.

56. Hall-Stoodley L, Hu FZ, Gieseke A, Nistico L, Nguyen D, Hayes J, et al. Direct detection of bacterial biofilms on the middle-ear mucosa of children with chronic otitis media. Jama. 2006;296(2):202–11.

57. Brook I. Microbiology of sinusitis. Proceedings of the American thoracic society. 2011;8(1):90–100.

